# TLmutation: predicting the effects of mutations using transfer learning

**DOI:** 10.1101/2020.01.07.897892

**Authors:** Zahra Shamsi, Matthew Chan, Diwakar Shukla

## Abstract

A reoccurring challenge in bioinformatics is predicting the phenotypic consequence of amino acid variation in proteins. With the recent advancements in sequencing techniques, sufficient genomic data has become available to train models that predict the evolutionary statistical energies, but there is still inadequate experimental data to directly predict functional effects. One approach to overcome this data scarcity is to apply transfer learning and train more models with available datasets. In this study, we propose a set of transfer learning algorithms we call TLmutation, which implements a supervised transfer learning algorithm that transfers knowledge from survival data of a protein to a particular function of that protein. This is followed by an unsupervised transfer learning algorithm that extends the knowledge to a homologous protein. We explore the application of our algorithms in three cases. First, we test the supervised transfer on 17 previously published deep mutagenesis datasets to complete and refine missing datapoints. We further investigate these datasets to identify which mutations build better predictors of variant functions. In the second case, we apply the algorithm to predict higher-order mutations solely from single point mutagenesis data. Finally, we perform the unsupervised transfer learning algorithm to predict mutational effects of homologous proteins from experimental datasets. These algorithms are generalized to transfer knowledge between Markov random field models. We show the benefit of our transfer learning algorithms to utilize informative deep mutational data and provide new insights into protein variant functions. As these algorithms are generalized to transfer knowledge between Markov random field models, we expect these algorithms to be applicable to other disciplines.

## Introduction

Proteins are intricate molecular machines that regulate all biological processes. The function of a particular protein is intrinsically linked to its structure, which governs its stability and conformational dynamics.^1,2^ Consequently, mutations in protein sequences, in which one amino acid is replaced by another amino acid, can affect a protein’s structure, stability, and inevitably its function. While some mutations may have little to no effect on a protein’s function, others have larger implications for disease and antibiotic resistance.^3,4^ Recent advancements in large-scale genomic sequencing have provided tools and resources for both consumers and clinicians to identify disease-potential mutations in one’s proteome at an affordable cost. However, due this large influx of genomic data, a recurring challenge is predicting the phenotypic consequence in proteins due to amino acid variations.^5,6^

Several workflows, both experimentally and computationally, have been developed to identify, predict, and model the effects of mutations.^7–10^ Engineering approaches, such as deep mutational scanning, provide a unique glimpse into the sequence-function relationship of proteins by surveying all single-point mutations in the sequence and assessing their altered function.^11–14^ These methods provide large quantitative datasets of mutational effects for a particular protein. Alternatively, statistical models have been used as standalone approaches or to compliment biophysical experiments. PolyPhen2^15^ and SIFT^16^ are examples of common frameworks that use multiple sequence and structural-based alignments to characterize variants. Other models, such as SNAP2,^17^ CADD,^18^ and Envision,^19^ employ machine learning algorithms to classify and predict mutations and are popular due to their robustness with large datasets. One successful approach employed for predicting mutational effects is EVmutation which uses evolutionary sequence conservation.^20^ In addition to applying evolutionary conservation to predict the effect of mutations, EVmutation also considers genetic interactions between mutations and the sequence background. By accounting for the interactions between all residue pairs, the model predicts the effects of mutations accurately as compared to other predictors. ^20^ Additionally, this method is shown to be able to capture the functionally relevant protein conformations and their dynamics.^21–24^ EVmutation utilizes a graph based Markov random field known as the Potts model which is trained on natural sequences.^20^ This means, for a given sequence, the algorithm searches through the UniProt database, ^25^ locates all natural sequences in its family, and uses these sequences as data to train the Potts model. However, these unsupervised probabilistic models do not directly predict the effects of mutations on the functionality of the protein; rather, they predict if the mutant species are fit to survive, which may not always directly correlate to a specific function. ^20^

As genomic sequencing and mutational libraries become readily available, it represents an opportunity to utilize this data-rich regime to enhance predictors of protein mutations. However, training Potts models on specific experimental data is usually not feasible. While mutational libraries of various proteins have been completed due to advancements in deep mutational scanning and directed evolution methods, ^26–28^ these datasets are inadequate for training, and we must rely on alternative methods to obtain insights and predictions. In a traditional machine learning approach, different task will be individually learned to build a model. However, in many real-world applications, collecting training data and rebuilding models may be computationally expensive.^29^ These difficulties are akin to deep mutational scanning and other biophysical approaches. Many experimental methods in characterizing protein variants are susceptible to noise or missing datapoints. ^9^ Moreover, it is difficult to infer information about other proteins from a single deep mutational scan.

One approach to overcome this data scarcity is to apply transfer learning algorithms in which we apply knowledge from one task to a different, yet related task. ^29^ Transfer learning has been used to tackle various challenges in molecular biology and bioinformatics, including protein function prediction,^30,31^ and protein-protein interactions.^32,33^ Singh *et al*. developed a platform to predict RNA secondary structure from models initially trained on a high-resolution RNA structure database. ^34^ Due to insufficient data on residue contacts in membrane proteins, Wang *et al*. transferred convolutional neural network parameters of non-membrane protein contacts to enhance structure prediction of membrane proteins. ^35^ While the motivation of employing machine learning is to enhance predictions in low data regime, transfer learning can take advantage of the structural and functional similarities between homologous proteins.

Here, we propose an algorithm, TLmutation, which is an adaptation of the successful variant effect predictor EVmutation, that utilizes deep mutational datasets to enhance predictions of variant effects in proteins. We implement the algorithm in two fashions. First, TLmutation transfers knowledge from a model, trained on natural sequences and deep mutational data, to a new protein function for the same protein. We call this algorithm supervised TLmutation. This is followed by an unsupervised transfer learning algorithm that expands the knowledge to a related protein and is referred to as unsupervised TLmutation. We conducted multiple experiments using the proposed transfer learning algorithms to evaluate the practical efficiency in predicting the effects of mutations of multiple proteins with different types of training and test datasets. In the first case, we explore the application of the proposed algorithm on 17 previously published mutagenesis datasets to complete missing datapoints. We further investigate different sampling approaches to delineate which mutations provide more accurate predictors of variant functions. In the second case, we apply our algorithm to predict higher order mutations (i.e. double mutations, triple mutations) solely from single point mutatgensis data. Finally, we implement the unsupervised transfer learning algorithm to predict mutational effects of homologous proteins from experimental datasets. Our results show that the incorporation of deep mutational dataset not only enhances the prediction of variant effects, but also can be transferable to provide new insights where experimental data may be limited.

## Methods

### Potts model for protein sequences

A Markov random field (MRF) is an undirected, probabilistic graphical model that represents statistical dependencies among a set of random variables, *σ* = (*σ*_1_,…, *σ_N_*), where ∀ *σ_i_* ∈ {1, 2,.., *l*}. MRF models have widely been used to tackle large datasets in different disciplines such as genomic biology,^36^ physics,^37^ natural language processing,^38^ and computer vision.^39^ For this study, let *σ* = (*σ*_1_, *σ*_2_,…, *σ_N_*) represent the amino acid sequence of a protein with length *N*. Each *σ_i_* takes on values in {1,2,…, 21} (one state for each of the 20 naturally occurring amino acids and one additional state to represent a gap). The probability of *σ*_1_,…, *σ_N_* is then given by:

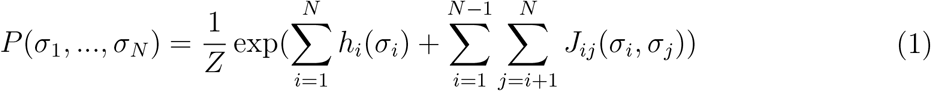

where *h_i_* is the potentials of site *i*, or fields, *J_ij_* is the potentials between residue pair constraints, or couplings of sites *i* and *j*,^20^ and *Z* is the partition function.^40^ This form of the MRF is commonly known as the Potts model or Potts Hamiltonian models.^41^

Assume we have two similar domains, source and target domain, in which we have a base and a new task (Figure 1). The new task must be a subset of the base task. This means the base task has to be the result of multiple smaller tasks, of which the new task is one of them. We define 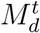 be a MRF *M* of the task *t* in the domain *d*. We initially are provided two MRF models for the source domain and the target domain, both trained on the base task, or 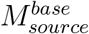 and 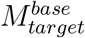, respectively. Our aim is to obtain MRF models on the new task for both source and target domains (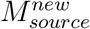 and 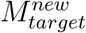). The training data for the source domain contains a set of *n* data points for the new task, 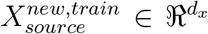, and its corresponding labels or outputs, 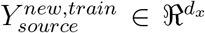. Additionally, we have test datasets for both the source and the target domain (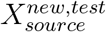 and 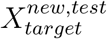, with labels as 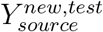 and 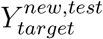, respectively).

**Figure 1:**
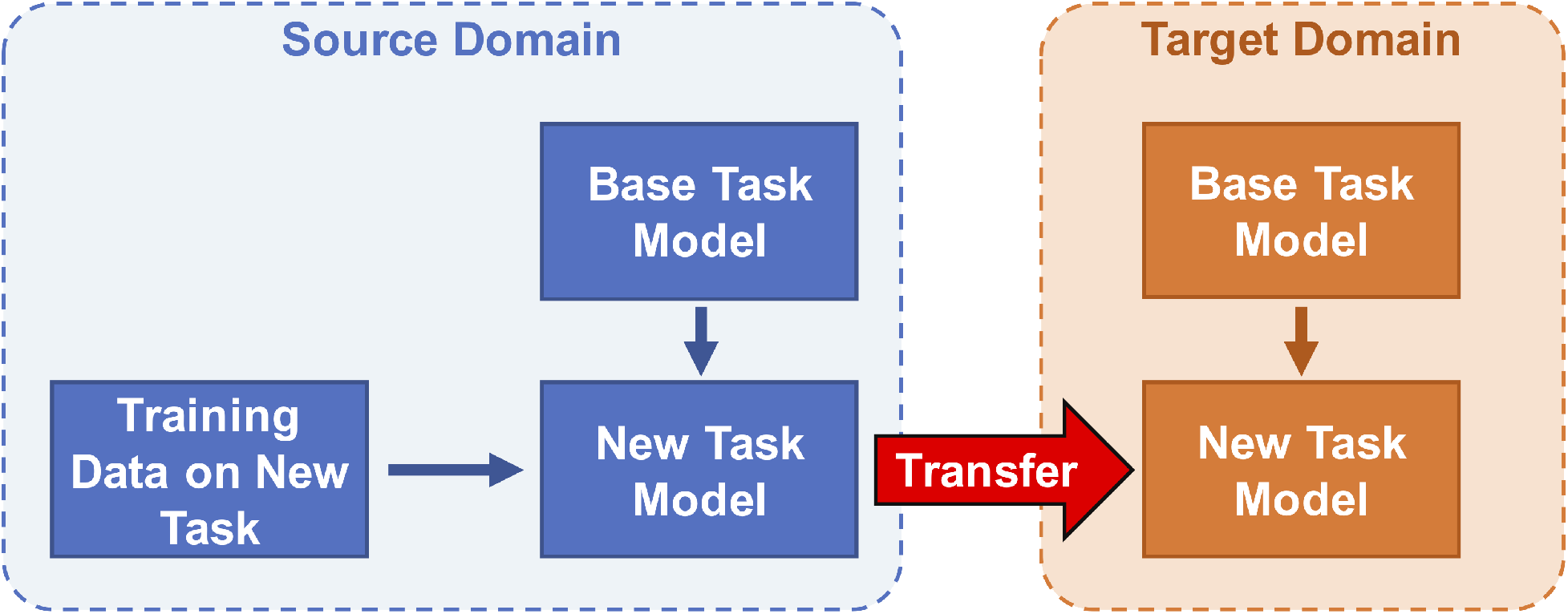
Proposed transfer learning algorithms for MRF models. In each source and target domain, the new task is a subset of the base task. Knowledge is then transferred from the source domain, where training data, is available to the target domain.

The first goal is to utilize the given training data and the MRF model to find a predictive model for the new task in the source domain 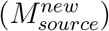. Then, we extend the knowledge from this supervised transfer learning step to learn a model in the target domain 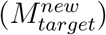 using an unsupervised transfer learning algorithm. To evaluate the performance of the algorithm, the predictions (*Ŷ*) are ranked and compared to the actual labels (*Y*) for calculation of the Spearman rank correlation coefficient (*ρ*).^42^ For this study, *ρ* accesses the association between two ranked variables, the predicted effect of a point mutation and the experimental effect. The value of *ρ* ranges from −1 to +1, where +1 indicates one variable is a perfect, monotonically increasing function of the other variable and −1 as a perfect, monotonically decreasing relationship.

### Supervised transfer from the evolutionary statistical energy to a functional assay

We want to modify an existing predictive model that is trained on the base task to be able to predict a new task. In the supervised transfer portion, all the transfer is conducted within a same domain, the source domain, and therefore the subscripts that determine the domains are dropped. First, we train a MRF model on the base task (*M^base^*) and calculate the values for the potentials of 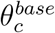 of all residues c using available methods in the literature.^20^ Then, we introduce a MRF model that can predict the new task as the following:

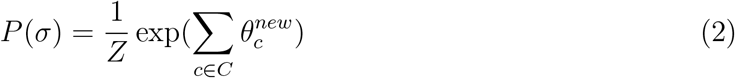

where

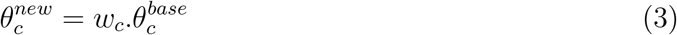

*w_c_* is a binary weight matrix (*w* ∈ {0, 1}), which is calculated by maximizing the correlation between the predicted values of labels (*Ŷ^new,train^*) and the actual labels (*Y^new,train^*) as

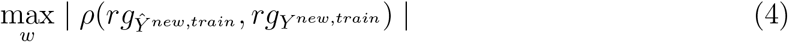

with *rg_Ŷ^new,train^_* and *rg_Y^new,train^_* are ranks of *Ŷ^new,train^* and *Y^new,train^*, respectively. *ρ* is the Spearman’s rank correlation coefficient. The maximization algorithm is further explained in the Supporting Information.

In this particular study, we want to learn a Potts model that can predict the effect of mutations on a particular protein function. We are given a Potts model trained on natural sequences (EVmutation model) and experimental data of the effects of point mutations on a protein’s function. In the Potts model of a protein sequence (Eq. 1), each *J_ij_* parameter represents the chemical or physical interactions between the corresponding residues *i* and *j*.^20^ Since the model is trained on survival data, *J_ij_*’s with large values suggest critical interactions necessary for survival. Such interactions may have roles including, but not limited to, expression, folding, thermal stability, or conformational dynamics. Assuming our function of interest is one of these essential survival functions, we want to decouple the *J_ij_* parameters from the overall survival by nullifying *J_ij_*’s that do not contribute to the function and retaining the ones that are linked to this function. This forms the basis of the proposed supervised transfer learning algorithm, supervised TLmutation.

In the supervised transfer learning section, all the transfer occurs within the same protein (referred to as source protein). First, we train a Potts model on survival (e.g. EVmutation), and calculate the values for the potentials of *J_ij_* and *h_i_*.^20^ Then, we introduce a new Potts model that can predict the function using the following modified potentials:

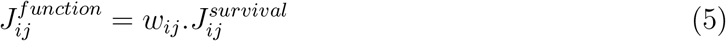

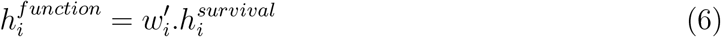

Analogous to the generalized modified potential (Eq. 2), *w_i.j_* and 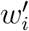 are binary weight matrices (*w_c_* ∈ {0, 1}) and is calculated by maximizing the correlation between the predicted values of the protein mutant’s function (*Ŷ^train^*) and the actual experimental values in the training set (*Y^train^*) (Eq. 4). The binary weights are used in the algorithm due to training data scarcity. The weight matrices for the couplings parameter *J_ij_* contains *n* * 21 *21 number of elements, where *n* is the length of the protein. Likewise, the weight vector for the fields parameter *h_i_* contains *n* * 21 number of elements. The total elements required to train the model is much more than the available experimental data points. Therefore, we decided to constrain the parameter state space using a binary mask. A similar use of the binary mask has recently been implemented for image recognition alogorithm. ^43^ Furthermore, we eliminate potentials which do not contribute to the enhancement in predicting the *Y^new,train^*. In this way, we eliminate the effects of other functions and focus on the function of interest.

### Unsupervised transfer between proteins

Now, we want to expand the knowledge gained from the supervised transfer to the target domain, where the training data is not available for the new task. Therefore, we train a MRF model on the base task in the target domain 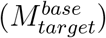 and a MRF model on the new task in the source domain 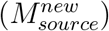. In our generalization example, using the learned model parameter, *w_c_*, from Equation 3, we define a MRF model for the new task in the target domain 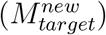. This model’s potential 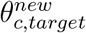 is calculated using Equation 7.

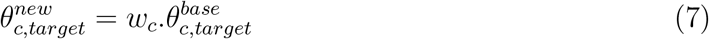

Since the source and target domains are similar, we assume the corresponding potentials have the same effects on predicting the new task. Therefore, we use the same learned weights *w* to switch the potentials in 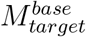 and obtain 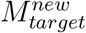 while using the same value of 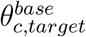 for potentials that remain active in the MRF.

Here, we extended the supervised and unsupervised TLmutation algorithms to a more generic MRF model. The algorithm transfers knowledge from a well trained MRF model of one task, to other similar tasks when either limited or no training data is available. The proposed supervised and unsupervised transfer algorithms presented can be generalized and applied to other MRF models. For most proteins, we do not have sufficient training data to use a supervised transfer learning algorithm as obtaining mutation data is experimentally challenging and expensive.^44,45^ We want to use the available experimental data of a protein for predicting the mutation effects in other homologous proteins. We expand the knowledge gained from the supervised TLmutation to the target protein, where no training data is available.

Assume the EVmutation model for target protein and the TLmutation model for source protein are constructed. Using the parameters of the TLmutation, *w* and *w′*, from Equation 5, we define a new Potts model for predicting the effects of mutations in the target protein on its function. The new model’s potentials are calculated using Equation 8 and 9.

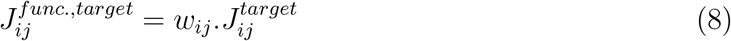

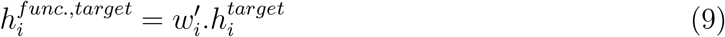

where *w_ij_* and 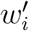 are the binary masks from the TLmutation model of the source protein. 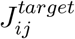 and 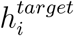 are the potentials from the EVmutation model of the target protein, and 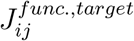 and 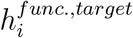 are the modified potentials for TLmutation model of the target protein. Here, the source and target proteins should be homologs, as the molecular mechanisms leading to a particular function are expected to be conserved among homologus proteins. The residue-residue interactions involved in the function of the source protein also are anticipated to be conserved in the target protein. Similarly, for the unsupervised transfer, we eliminate the *J_ij_*’s in the target protein that did not contribute to the function in the source protein. However, we used the target’s EVmutation model as the preliminary basis and applied the binary masks from source to its potentials.

## Results

### Case study 1: Filling gaps in experimental datasets

In this first case study, we want to evaluate the performance of TLmutation in refining and filling gaps in the deep mutagenesis datasets. It is common to have incomplete mutagenesis datasets, where readings of certain variants were unobtainable due to experiemntal difficulties (e.g. expression or purification of protein, poor sequencing). Here, we apply the TLmutation algorithm to predict missing variants of 17 previously published, large-scale mutagensises datasets (see SI Table S1 for more details). These datasets include quantitative measurements of variant effects for various protein functions. For each dataset, the available data were randomly divided into test and training sets. 5-fold cross-validation was used to assess the robustness of the supervised TLmutation models. Our initial analysis showed TLmutation does not significantly improve EVmutation models for incomplete datasets which have less than 6 mutations per site of the mutagenized region. This leads us to enforce an additional constraint on the datasets to include experimental values for at least ~ 35% of all possible variants of the mutagenized region (Figure S1). As shown in Figure 2 **B**, 12 out of 17 datasets contain an adequate numbers of experimental datapoints. In these datasets, supervised TLmutation improved the correlation coefficient with the actual experimental values for the test set in 12 of the 13 proteins (p-value < 0.05) (Figure 2 **C**). Among these, the largest improvements are observed in systems with a lower correlation between EVmutation and experiments.

**Figure 2:**
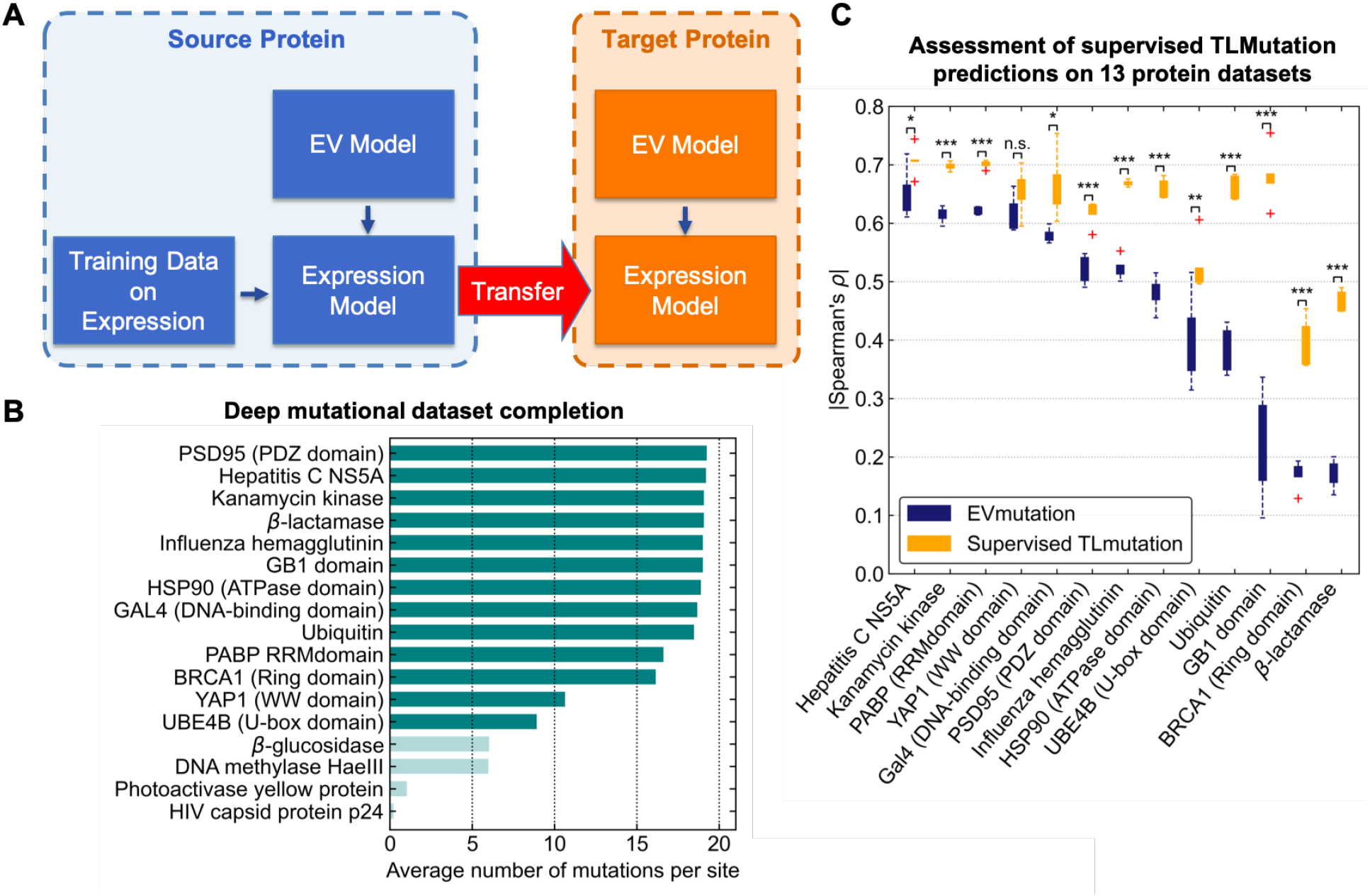
(**A**) Schematic of the proposed transfer learning algorithms on an example of predicting protein expression. Knowledge is transferred from the source protein where training data is available to the target protein. (**B**) Completeness of 17 studied mutagenesis datasets is shown. Datasets shown in light teal have less than 35% of all possible variants in the mutagenized region. (**C**) Effects of mutations computed using EVmutation and supervised TLmutation. These predictions are compared with experimental measurements for 13 proteins are shown for the test set. The agreement is measured by Spearman’s rank correlation coefficient *ρ*. Asterisks indicate statistically significant (*, *p* < 0.05; **, *p* < 0.01; ***, *p* < 0.001, *n.s*., not significant). Error bars are calculated from 5-fold cross-validation. Outliers are represented as red crosses.

#### Which mutations should be experimentally tested to build better predictors of variant function?

As there are more than thousands of possible sites on a protein that can be subjected to mutagensis, it is beneficial to understand which datapoints may provide the most generalizable information about other variants. Here, we address this question by training the TLmutation model using different sampling methods.

Simple random sampling is the most common method as it is efficient and relatively easy to implement. In this approach, samples are randomly selected with a uniform distribution. This sampling method was used in the previous section and showed a significant improvement in the performance of the model in all datasets (Figure 2 **C**, training scores are shown in Figure S2). However, mutagenesis datasets are inherently not uniform. More sophisticated sampling methods will be more suitable for these types of naturally ordered datasets. Here, we tested two systematic sampling approaches on the same 13 protein datasets. In the first approach, we divided the datasets based on sequence or positions of the proteins. The TLmutation model was trained on all available mutations for 80% of the sequence sites and tested on the remaining data (as shown in Figure 3 **A**). As before, 5-fold cross-validation was used to assess the sampling method. This approach dramatically decreased the performance of the model. In most of the systems, no improvement was observed as compared to EVmutation (Figure 3 **B**). This observation suggests that the effects of mutations on sites far away from each other may not correlate with the mutations in other sites. This leads us to the second sampling approach, where the test/training splitting occurred for each position, meaning that 80% of available experiments for mutations on each sequence site was labeled as training, and 20% as test (as shown in Figure 3 **C**). Using this sampling, TLmutation outperformed EVmutation in 12 of the 13 systems (Figure 3 **D**). However, the improvement is still comparable with random sampling.

**Figure 3:**
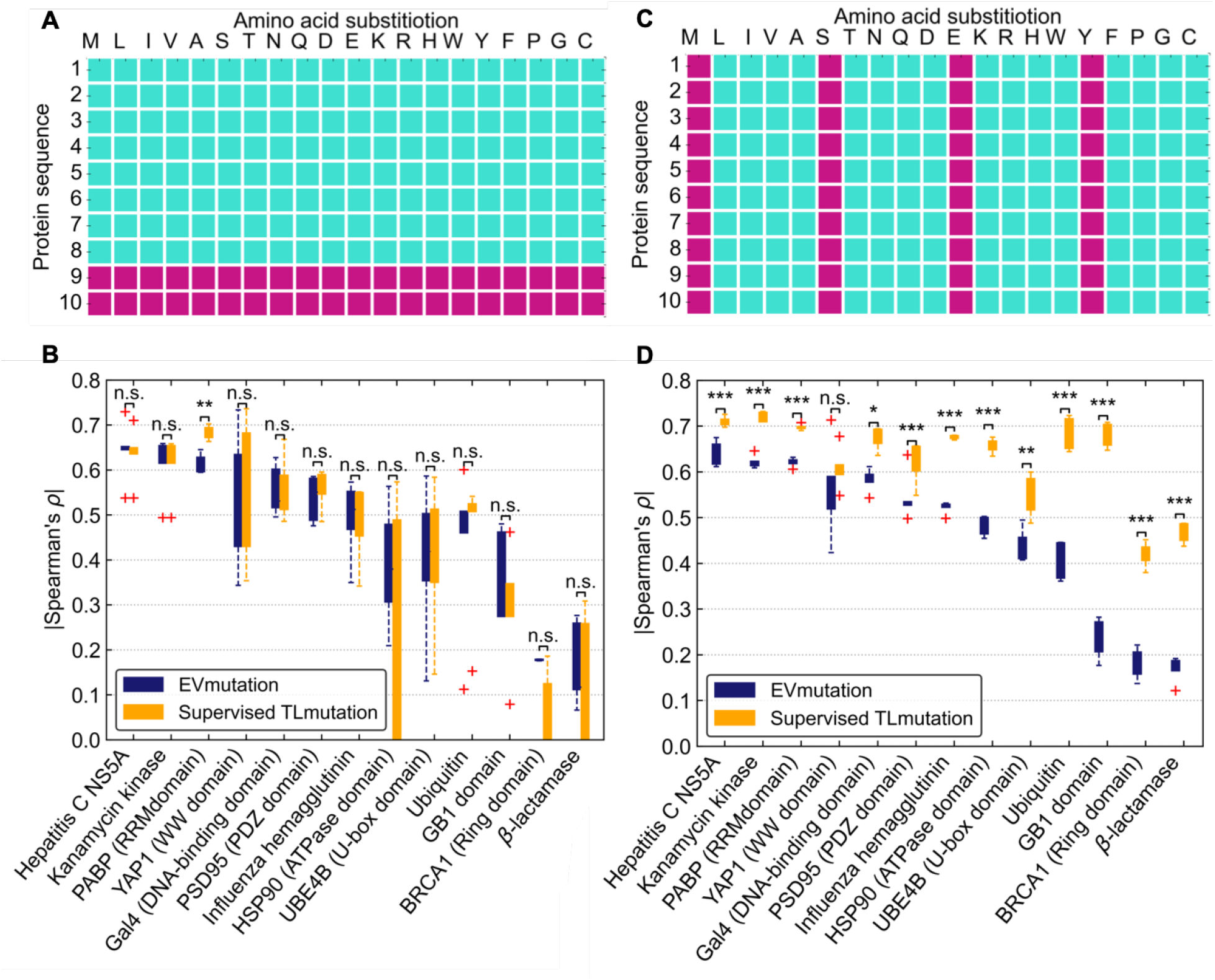
(**A**) A demonstration of train/test split based on sequence positions is shown. Purple points are considered in the test set and teal points in the training set. (**B**) The correlation between computed effects of mutations and experimental measurements for 13 proteins are shown for 5-fold cross validation and site based train/test split. Outliers shown as red crosses. TLmutation using site based train/test split, does not significantly improve the correlation coefficients. Asterisks indicate statistically significant (*, *p* < 0.05; **, *p* < 0.01; ***, *p* < 0.001, *n.s*., not significant). (**C**) A demonstration of train/test split based on amino acid substitutions is shown. (**D**) The correlation between computed effects of mutations and experimental measurements for 12 proteins are shown for 5-fold cross validation and substituted amino acid based train/test split. TLmutation improves the correlation coefficients in 12 of the 13 datasets. Asterisks indicate statistically significant (*, *p* < 0.05; **, *p* < 0.01; ***, *p* < 0.001, n.s., not significant).

### Case study 2: Predicting the effects of multiple point mutations from single point mutation experiments

While a single point mutation may not affect the protein’s function, it is possible for multiple mutations to cooperatively affect its function. Datasets of single point mutations have become increasing available over the past decade. However, mutational maps with multiple point mutations remains scare and are difficult to obtain experimentally. From an experimental perspective, the number of possible mutants increases exponentially with the increase in number of mutated residues, thus conducting a thorough mutagenesis analysis of large proteins is challenging and expensive. Here, we use the supervised TLmutation algorithm to train on the available single point mutations and predict the effects of multiple point mutations. We tested the performance of the algorithm on 4 previously published mutagenesis datasets. These datasets have been employed in the literature to evaluate the performance of EVmutation. ^20^ These datasets contain single and double point mutations (more detail is provided in SI table S2). In these systems, the correlation coefficient was increased for both test and training sets as compared to EVmutation (*p*-value < 0.05) (Figure 4). Our results show that the incorporation of experimental mutant data combined with couplings derived from the model allows accurate predictions of the effects of higher order mutations. This use of deep mutational data and couplings has also been leveraged to predict three-dimensional structures of proteins.^46,47^

**Figure 4:**
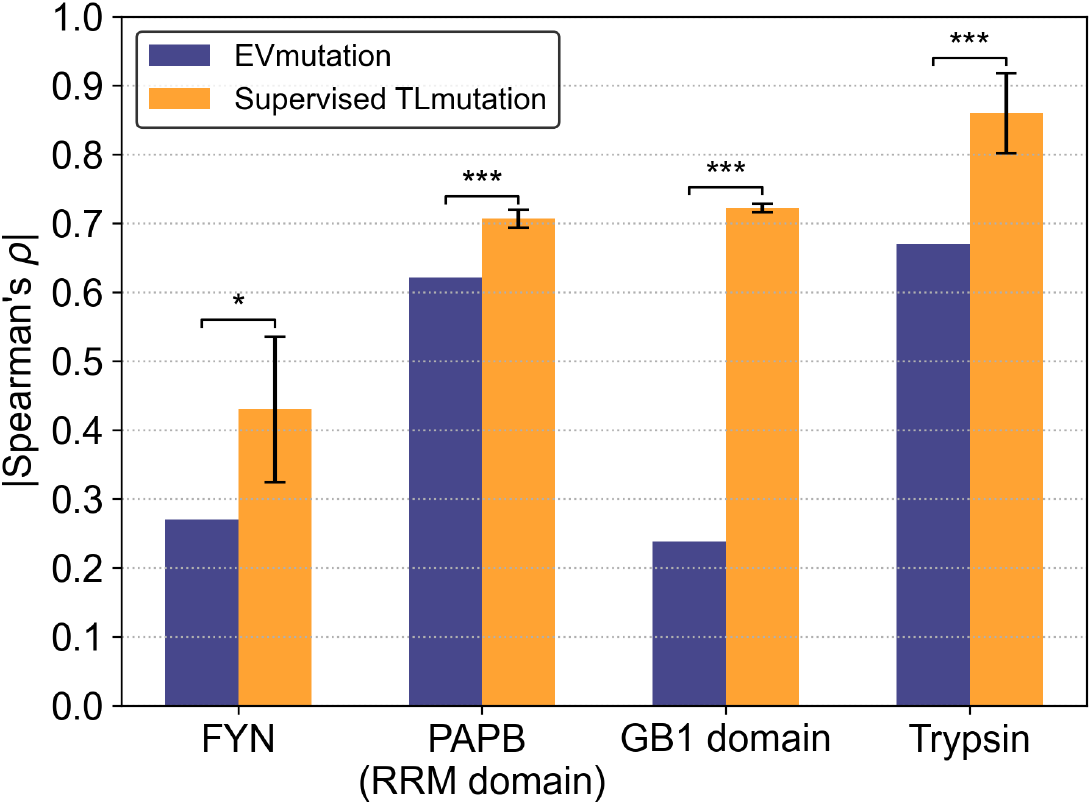
The agreement between predicted effects of double point mutations with experimental measurements is shown for the test datasets. The supervised TLmutation models were first trained on limited available single point mutation data and used to predict double mutation effects. Incorporation of experimental data significantly enhances prediction of double mutants. Asterisks indicate statistically significant (*, *p* < 0.05; ***, *p* < 0.001).

### Case study 3: Predicting the effects of mutations using available experimental data on a homologous protein

Mutagensis datasets provide an opportunity to utilize this data-rich regime and investigate the transferability of mutational effects among homologous proteins. The effectiveness of our transfer algorithms was tested for two chemokine receptors, CXCR4 and CCR5 (Figure 5). Chemokine receptors belong to the class of G-protein coupled receptors (GPCRs) that transmit cellular signals across the cell membrane on the binding of signaling molecules known as chemokines on the extracellular side.^48^ These receptors regulate the movement of immune cells in the body, most notably white blood cells during inflammation.^49^ Chemokine receptors play a vital role in HIV-1 infection and progression,^50^ and hence are considered as major drug targets for treating HIV-1, along with other autoimmune disorders and cancer. ^51,52^ Specifically, both CXCR4 and CCR5 have been identified as co-receptors for HIV-1 entry into immune cells. Numerous efforts have been made to understand HIV pathology and to develop new therapeutic approaches. Clinical studies have associated a lack or low expression of CCR5 to provide a natural resistance of HIV infection.^53^ Mutational analysis, for example the deep mutational scanning of these receptors could provide an invaluable insights into the function of these receptors albiet at a high experimental cost. ^11,45^ There are more than 20 chemokine receptors in the human body, ^54^ of which only 2 currently have a mostly complete mutational dataset. ^55^ Evaluating the TLmutation algorithms on these two datasets would allow us to uncover the sequence-function relationship for other chemokine receptors.

**Figure 5:**
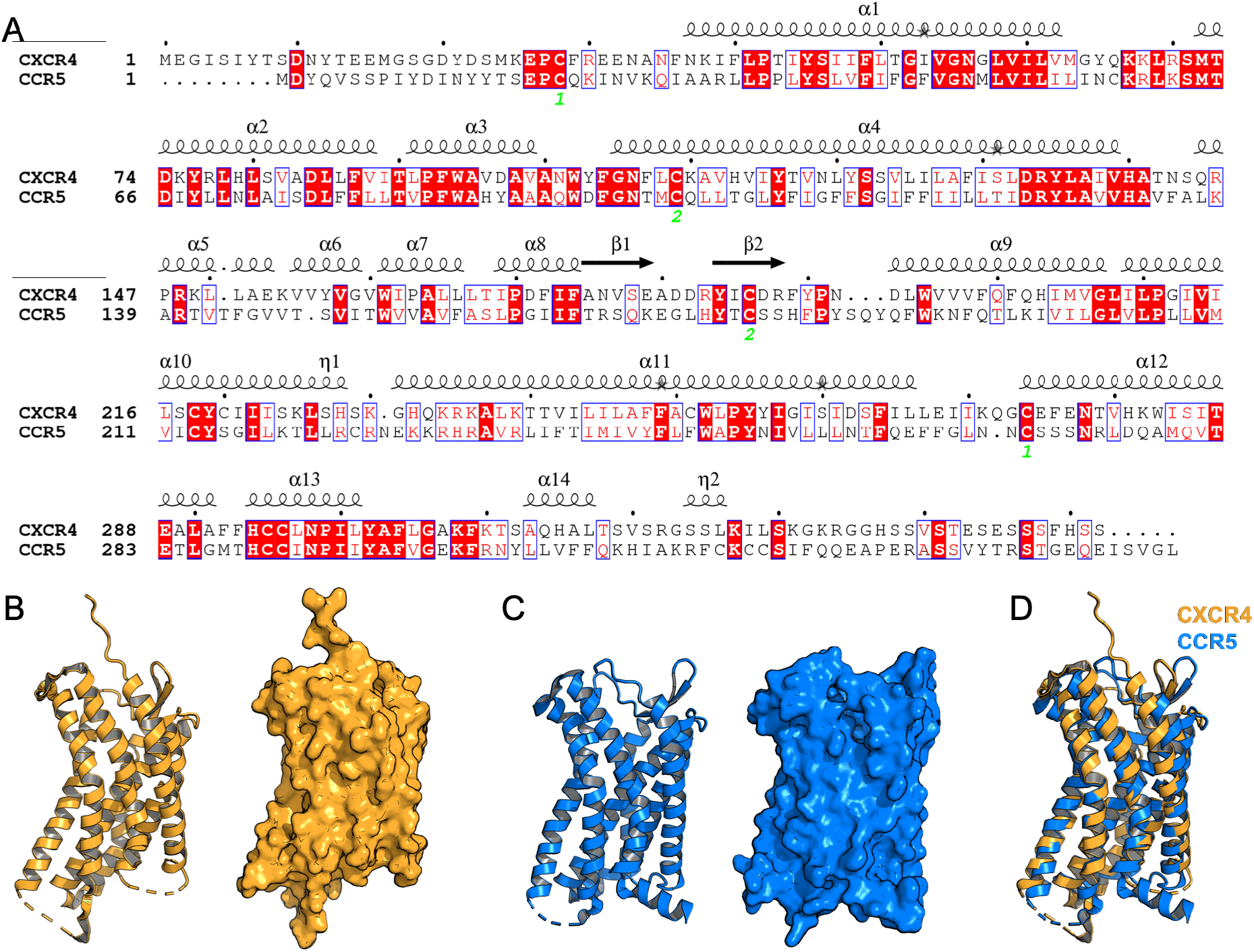
(**A**) Sequence alignments of chemokine receptors CXCR4 and CCR5. Identical residues colored in red boxes, where similar residues are colored in red letters. Corresponding secondary structure is placed above the aligned sequence. The amino acid sequences of the two receptors share a 30% sequence identity and 50% sequence similarity. Crystal structures, depicted in cartoon and surface representation, of CXCR4 (PDB: 4RWS) (**B**) and CCR5 (PDB: 4MBS) (**C**). (**D**) Cartoon representation of CXCR4 superimposed on CCR5 shows the high structural similarity between these receptors.

In this case study, we utilized available single point mutation datasets for two proteins, CXCR4 and CCR5, and two different experiments, expression level and bimolecular fluorescence complementation (BiFC) assay.^55^ Sequence identity and similarity between CXCR4 and CCR5 are ~ 30% and 50%, respectively (Figure 5). The proposed supervised and unsupervised TLmutation algorithms were implemented and tested for one case of supervised transfer and four different cases of unsupervised transfer. The EVmutation models for CCR5 and CXCR4 were built using 95619 and 94461 natural sequences using the algorithm as explained in the literature to compare its performance with the TLmutation.^20^

We want to learn a Potts model that can predict the effects of mutations on expression levels of CXCR4 given a Potts model trained on natural sequences related to CXCR4 and its expression levels for 6994 single point CXCR4 mutants. Using the supervised TLmutation algorithm, a 5-fold cross-validation was performed for the available experimental data on expression levels of CXCR4. The Spearman coefficients for the training data increased from 0.174±0.006 to 0.403±0.005 and the coefficients for the test data increased from 0.174±0.023 to 0.279 ± 0.041 as it is shown in Figure 6 **C**. For all of the 5 folds, the correlation between predicted ranks and actual experiment is higher compared to EV model. For fold 3, the projection of the combined error for each residue is shown on the 3D structure of CXCR4 in Figure 6 **A** and **B**. The overall error is considerably lower for the proposed algorithm as compared to EVmutation model. ^20^

**Figure 6:**
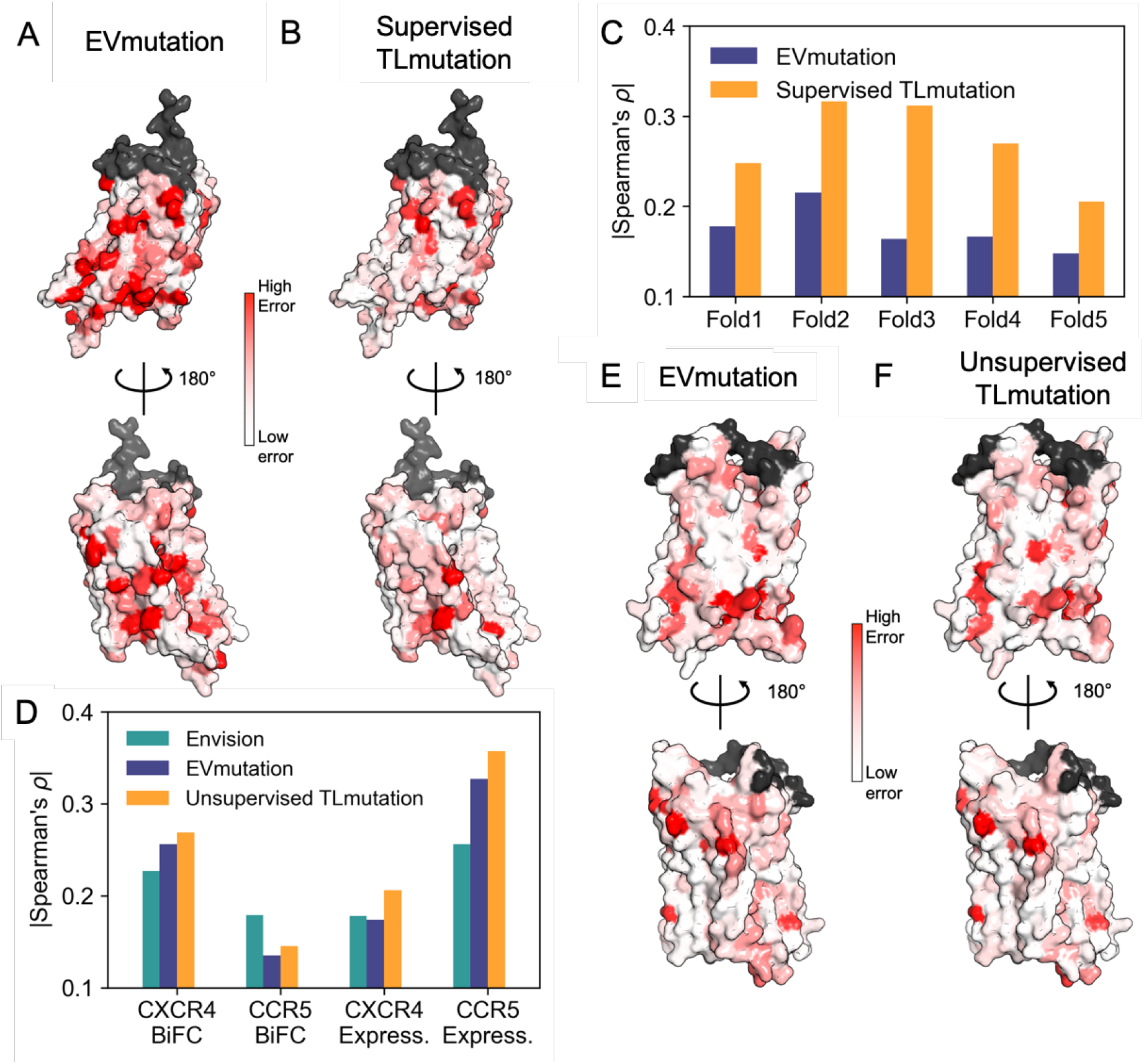
Comparison of the CXCR4 expression levels predicted from the EVmutation (**A**) and supervised TLmutation (**B**). Color indicates relative error with respect to experimental value, ranging from low (white) to high (red). Residues of low-confidence predictions (<20 residues) are colored in grey. (**C**) Test Spearman coefficients in 5-fold cross-validation of CXCR4 on predicting the expression levels. Correlation with experimental values is shown on the Y axis, which measures the performance of the predictive model. Higher Spearman coefficient indicates better match with the data. (**D**) Performance of unsupervised TLmutation in regards to predicting different experiments for the target protein is compared with EVmutation and Envision. For these cases, training data sets are unavailable. The knowledge is transferred to the target in order to predict mutagenesis effects. Error in prediction of CCR5 expression as compared to experimental data using (**E**) EVmutation and (**F**) unsupervised TLmutation algorithm.

The effectiveness of the unsupervised TLmutation algorithm in predicting expression levels and BiFC was evaluated on four different cases: (1) transfer from CXCR4 to CCR5 expression, (2) CXCR4 to CCR5 BiFC, (3) CCR5 to CXCR4 expression, and (4) CCR5 to CXCR4 BiFC, as shown in Table 1 and Figure 6 **D**. Spearman coefficients in the third and fourth columns in Table 1 illustrate the improved performance of the supervised transfer on the training data compared to the EV model and Envision. The sixth and seventh columns indicate the Spearman coefficients of test data for EVmutation model and using the unsupervised TL, respectively. In all the cases, we observed that the proposed approach shows improvement over the current genomic prediction method (EVmutation model). For the first case (the first row of Table 1), where the supervised transfer is performed on expression levels of CXCR4, and the unsupervised transfer is to CCR5, we project the prediction errors of each residue onto the crystal structure (Figure 6 **E** and **F**). Likewise, the prediction errors are lower for the proposed unsupervised transfer, as compared to the EVmutation model.

**Table 1:** Unsupervised transfer learning between homologous chemokine receptors. Higher Spearman coefficient indicates better match with the actual data. Correlations are an average of 5 replicates obtained by using 90% of the source domain, randomly selected, as the training set followed by prediction of the effect of mutations in the target domain.

| Source Protein | New Task (Experiment) | EV $\rho$ (for Source) | Supervised TL $\rho$ (for Source) | Target Protein | EV $\rho$ (for Target) | Unsupervised TL $\rho$ (for Target) |
| --- | --- | --- | --- | --- | --- | --- |
| CCR5 | BiFC | 0.134±0.006 | 0.353±0.003 | CXCR4 | 0.256 | 0.269±0.003 |
| CXCR4 | BiFC | 0.257±0.005 | 0.457±0.005 | CCR5 | 0.135 | 0.146±0.001 |
| CCR5 | Expression | 0.327±0.002 | 0.518±0.004 | CXCR4 | 0.174 | 0.207±0.001 |
| CXCR4 | Expression | 0.175±0.004 | 0.397±0.003 | CCR5 | 0.326 | 0.367±0.002 |

## Discussion

In this work, we aimed to extract new predictive models of the biological function of proteins from predictive models trained on the evolutionary data and extending it to a new protein via unsupervised transfer learning. We showed that these techniques are efficient for multiple case studies, and compared the results with successful variant predictors EVmutation^20^ and Envision. ^19^ Another aspect of the work was focused toward understanding which subsets of data points yielded the most informative predictions. We showed, through our algorithm, that having few single point mutations on each position was sufficient to estimate the effects of other point mutations in the same sites. As these experiments are challenging and the data points may not contain uniform information, one natural followup of our work is to extend the algorithm to an active learning approach. In this scheme, we can suggest which mutations should be tested over few rounds and adaptively strengthen the model. Additionally, as these datasets are susceptible to noise, one approach is to implement the algorithm to enhance low-confidence predictions. By obtaining training data on the parts of the dataset which provide maximum information gained, we can reduce the number of experiments and train more powerful models with limited amounts of data.

One of the questions that remains unanswered in this work, is how to define the degree of transferability between domains and tasks. In this application, we compared the proteins based on their sequence and structural similarities. However as we do not have enough datasets to check the transferability between multiple proteins, it is challenging to define criterion that would allow successful transfer of knowledge between similar proteins. The example in this paper explored the transferability between CXCR4 and CCR5 proteins with ~ 30% sequence identity. When decoupling the survival parameters that do not contribute to a particular function of the protein, we assume that the experimental assay from deep mutagenesis is a measure of this function. However, this may not always be the case and may result in low correlations or differences in correlations between experiments. Additionally, while the functional residues between homologous protein may be similar, we do not account for the differences in the mechanism of these functions that are not represented in the model. For example, Sultan et al.^56^ have shown that the a neural network model trained for a protein using molecular dynamics trajectory data can efficiently be transferred to perform enhanced conformational sampling on a related mutant protein. However, Moffett and Shukla have shown that the transferrability of functional dynamics between related proteins is not always high and it depends on the differences between the functional free energy landscapes of protein and its mutants. ^57^ Therefore, while we expect systems of high sequence identity or structural similarity to be more transferable, we cannot validate this claim due to the lack of mutational datasets of homologous proteins.

Overall, we anticipate supervised TLmutation will be useful in predicting the effects of multiple point mutations and filling out gaps in mutagenesis datasets. Unsupervised TLmutation will help to expand the knowledge to predict the effects of the mutation in many homologous proteins. We expect unsupervised TLmutation to continue to improve as more datasets of homologous proteins become available. Furthermore, the proposed transfer learning algorithms were shown to be generalizable to all MRF models, which could be applicable to other disciplines.

## Supporting information

Supplementary Methods and Results

## Acknowledgement

The authors thank the Blue Waters sustained-petascale computing project, which is supported by the National Science Foundation (awards OCI-0725070 and ACI-1238993) and the state of Illinois. Blue Waters is a joint effort of the University of Illinois at Urbana-Champaign and its National Center for Supercomputing Applications. DS acknowledges support from the NSF Early Career Award, NSF MCB-1845606. All the code needed to generate the models presented in this work are available freely online under the MIT license at https://github.com/ShuklaGroup/TLMutation.

## Graphical TOC Entry

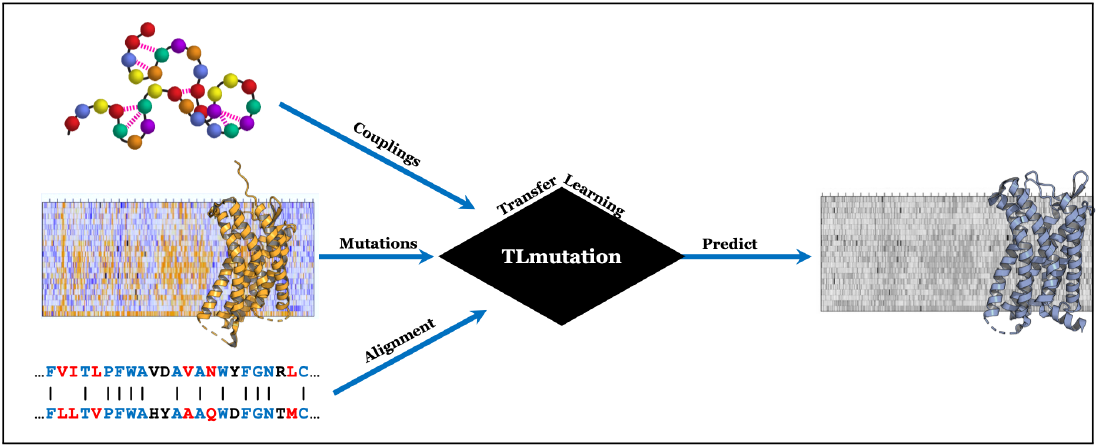

## References

(1) Henzler-Wildman, K.; Kern, D. Dynamic personalities of proteins. Nature 2007, 450, 964–972.

(2) Shukla, D.; Hernández, C. X.; Weber, J. K.; Pande, V. S. Markov State Models Provide Insights into Dynamic Modulation of Protein Function. Acc. Chem. Res. 2015, 48, 414–422.

(3) Wang, Z.; Moult, J. SNPs, protein structure, and disease. Hum. Mutat. 2001, 17, 263–270.

(4) Martinez, J.; Baquero, F. Mutation frequencies and antibiotic resistance. Antimicrob. Agents Chemother 2000, 44, 1771–1777.

(5) Shendure, J.; Akey, J. M. The origins, determinants, and consequences of human mutations. Science 2015, 349, 1478–1483.

(6) Mann, J. K.; Barton, J. P.; Ferguson, A. L.; Omarjee, S.; Walker, B. D.; Chakraborty, A.; Ndung’u, T. The fitness landscape of HIV-1 gag: advanced modeling approaches and validation of model predictions by in vitro testing. PLoS Comput. Biol. 2014, 10, e1003776.

(7) Gray, V. E.; Hause, R. J.; Fowler, D. M. Analysis of large-scale mutagenesis data to assess the impact of single amino acid substitutions. Genetics 2017, 207, 53–61.

(8) Stein, A.; Fowler, D. M.; Hartmann-Petersen, R.; Lindorff-Larsen, K. Biophysical and mechanistic models for disease-causing protein variants. Trends Biochem. Sci. 2019, 44, 575–588.

(9) Fowler, D. M.; Stephany, J. J.; Fields, S. Measuring the activity of protein variants on a large scale using deep mutational scanning. Nat. Protoc. 2014, 9, 2267.

(10) Riesselman, A. J.; Ingraham, J. B.; Marks, D. S. Deep generative models of genetic variation capture the effects of mutations. Nat. Methods 2018, 15, 816–822.

(11) Fowler, D. M.; Fields, S. Deep mutational scanning: a new style of protein science. Nat. Methods 2014, 11, 801.

(12) Matreyek, K. A.; Stephany, J. J.; Fowler, D. M. A platform for functional assessment of large variant libraries in mammalian cells. Nucleic Acids Res. 2017, 45, e102–e102.

(13) Park, J.; Selvam, B.; Sanematsu, K.; Shigemura, N.; Shukla, D.; Procko, E. Structural architecture of a dimeric class C GPCR based on co-trafficking of sweet taste receptor subunits. J. Biol. Chem. 2019, 294, 4759–4774.

(14) Harris, D. T.; Wang, N.; Riley, T. P.; Anderson, S. D.; Singh, N. K.; Procko, E.; Baker, B. M.; Kranz, D. M. Deep Mutational Scans as a Guide to Engineering High Affinity T Cell Receptor Interactions with Peptide-bound Major Histocompatibility Complex. J. Biol. Chem. 2016, 291, 24566–24578.

(15) Adzhubei, I. A.; Schmidt, S.; Peshkin, L.; Ramensky, V. E.; Gerasimova, A.; Bork, P.; Kondrashov, A. S.; Sunyaev, S. R. A method and server for predicting damaging missense mutations. Nat. Methods 2010, 7, 248.

(16) Ng, P. C.; Henikoff, S. SIFT: Predicting amino acid changes that affect protein function. Nucleic Acids Res. 2003, 31, 3812–3814.

(17) Hecht, M.; Bromberg, Y.; Rost, B. Better prediction of functional effects for sequence variants. BMC Genomics 2015, 16, S1.

(18) Kircher, M.; Witten, D. M.; Jain, P.; O’roak, B. J.; Cooper, G. M.; Shendure, J. A general framework for estimating the relative pathogenicity of human genetic variants. Nat. Genet. 2014, 46, 310.

(19) Gray, V. E.; Hause, R. J.; Luebeck, J.; Shendure, J.; Fowler, D. M. Quantitative missense variant effect prediction using large-scale mutagenesis data. Cell Syst. 2018, 6, 116–124.

(20) Hopf, T. A.; Ingraham, J. B.; Poelwijk, F. J.; Schärfe, C. P.; Springer, M.; Sander, C.; Marks, D. S. Mutation effects predicted from sequence co-variation. Nat. Biotechnol. 2017, 35, 128.

(21) Shamsi, Z.; Moffett, A. S.; Shukla, D. Enhanced unbiased sampling of protein dynamics using evolutionary coupling information. Sci. Rep. 2017, 7, 12700.

(22) Feng, J.; Shukla, D. Characterizing Conformational Dynamics of Proteins Using Evolutionary Couplings. J. Phys. Chem. B 2018, 122, 1017–1025.

(23) Feng, J.; Shukla, D. FingerprintContacts: Predicting Alternative Conformations of Proteins from Coevolution. J. Phys. Chem. B 2020, doi: 10.1021/acs.jpcb.9b11869.

(24) Shamsi, Z.; Cheng, K. J.; Shukla, D. Reinforcement Learning Based Adaptive Sampling: REAPing Rewards by Exploring Protein Conformational Landscapes. J. Phys. Chem. B 2018, 122, 8386–8395.

(25) Berman, H. M.; Westbrook, J.; Feng, Z.; Gilliland, G.; Bhat, T. N.; Weissig, H.; Shindyalov, I. N.; Bourne, P. E. The protein data bank. Nucleic Acids Res. 2000, 28, 235–242.

(26) Starita, L. M.; Young, D. L.; Islam, M.; Kitzman, J. O.; Gullingsrud, J.; Hause, R. J.; Fowler, D. M.; Parvin, J. D.; Shendure, J.; Fields, S. Massively parallel functional analysis of BRCA1 RING domain variants. Genetics 2015, 200, 413–422.

(27) Starita, L. M.; Pruneda, J. N.; Lo, R. S.; Fowler, D. M.; Kim, H. J.; Hiatt, J. B.; Shendure, J.; Brzovic, P. S.; Fields, S.; Klevit, R. E. Activity-enhancing mutations in an E3 ubiquitin ligase identified by high-throughput mutagenesis. Proc. Natl. Acad. Sci. U.S.A. 2013, 110, E1263–E1272.

(28) Heredia, J. D.; Park, J.; Choi, H.; Gill, K. S.; Procko, E. Conformational engineering of HIV-1 Env based on mutational tolerance in the CD4 and PG16 bound states. J. Virol. 2019, 93, e00219–19.

(29) Pan, S. J.; Yang, Q. A survey on transfer learning. IEEE Transactions on knowledge and data engineering 2009, 22, 1345–1359.

(30) Nauman, M.; Rehman, H. U.; Politano, G.; Benso, A. Beyond homology transfer: deep learning for automated annotation of proteins. J. Grid Comput. 2019, 17, 225–237.

(31) Rao, R.; Bhattacharya, N.; Thomas, N.; Duan, Y.; Chen, P.; Canny, J.; Abbeel, P.; Song, Y. Evaluating protein transfer learning with TAPE. Advances in Neural Information Processing Systems. 2019; pp 9686–9698.

(32) Mei, S. Probability weighted ensemble transfer learning for predicting interactions between HIV-1 and human proteins. PLOS ONE 2013, 8.

(33) Qi, Y.; Tastan, O.; Carbonell, J. G.; Klein-Seetharaman, J.; Weston, J. Semi-supervised multi-task learning for predicting interactions between HIV-1 and human proteins. Bioinformatics 2010, 26, i645–i652.

(34) Singh, J.; Hanson, J.; Paliwal, K.; Zhou, Y. RNA secondary structure prediction using an ensemble of two-dimensional deep neural networks and transfer learning. Nat. Commun. 2019, 10, 1–13.

(35) Wang, S.; Li, Z.; Yu, Y.; Xu, J. Folding membrane proteins by deep transfer learning. Cell Syst. 2017, 5, 202–211.

(36) Wei, Z.; Li, H. A Markov random field model for network-based analysis of genomic data. Bioinformatics 2007, 23, 1537–1544.

(37) Chen, W.; Jin, X.; Li, Z.; Zhang, X.; Hong, L. Clock Synchronization for Distributed Multi-hop Wireless Networks Using Markov Random Field. J. Phys. Conf. Ser. 2018; p 052008.

(38) Jernite, Y.; Rush, A.; Sontag, D. A fast variational approach for learning Markov random field language models. International Conference on Machine Learning. 2015; pp 2209–2217.

(39) Li, S. Z. Markov random field models in computer vision. Comput. Vis. ECCV. 1994; pp 361–370.

(40) Kindermann, R.; Snell, J. L. Markov Random Fields and Their Applications; American Mathematical Society., 1980.

(41) Levy, R. M.; Haldane, A.; Flynn, W. F. Potts Hamiltonian models of protein covariation, free energy landscapes, and evolutionary fitness. Curr. Opin. Struc. Biol. 2017, 43, 55–62.

(42) Myers, L.; Sirois, M. J. Spearman correlation coefficients, differences between. Encyclopedia of statistical sciences 2004, 12.

(43) Mallya, A.; Davis, D.; Lazebnik, S. Piggyback: Adapting a single network to multiple tasks by learning to mask weights. Proceedings of the European Conference on Computer Vision (ECCV). 2018; pp 67–82.

(44) Zhang, W.; Dourado, D. F.; Fernandes, P. A.; Ramos, M. J.; Mannervik, B. Multidimensional epistasis and fitness landscapes in enzyme evolution. Biochem J. 2012, 445, 39–46.

(45) Jonson, P. H.; Petersen, S. B. A critical view on conservative mutations. Protein Eng. 2001, 14, 397–402.

(46) Rollins, N. J.; Brock, K. P.; Poelwijk, F. J.; Stiffler, M. A.; Gauthier, N. P.; Sander, C.; Marks, D. S. Inferring protein 3D structure from deep mutation scans. Nat. Genet. 2019, 51, 1170.

(47) Schmiedel, J. M.; Lehner, B. Determining protein structures using deep mutagenesis. Nat. Genet. 2019, 51, 1177.

(48) Proudfoot, A. E. Chemokine receptors: multifaceted therapeutic targets. Nat. Rev. Immunol. 2002, 2, 106.

(49) Hughes, C. E.; Nibbs, R. J. A guide to chemokines and their receptors. FEBS J. 2018, 285, 2944–2971.

(50) Wang, Z.; Shang, H.; Jiang, Y. Chemokines and Chemokine Receptors: Accomplices for Human immunodeficiency virus infection and Latency. Front. Immunol. 2017, 8, 1274.

(51) Weitzenfeld, P.; Ben-Baruch, A. The chemokine system, and its CCR5 and CXCR4 receptors, as potential targets for personalized therapy in cancer. Cancer Lett. 2014, 352, 36–53.

(52) Chow, M. T.; Luster, A. D. Chemokines in cancer. Cancer Immunol. Res. 2014, 2, 1125–1131.

(53) Ostrowski, M. A.; Justement, S. J.; Catanzaro, A.; Hallahan, C. A.; Ehler, L. A.; Mizell, S. B.; Kumar, P. N.; Mican, J. A.; Chun, T.-W.; Fauci, A. S. Expression of chemokine receptors CXCR4 and CCR5 in HIV-1-infected and uninfected individuals. J. Immunol. Res. 1998, 161, 3195–3201.

(54) Griffith, J. W.; Sokol, C. L.; Luster, A. D. Chemokines and chemokine receptors: positioning cells for host defense and immunity. Annu. Rev. Immunol. 2014, 32, 659–702.

(55) Heredia, J. D.; Park, J.; Brubaker, R. J.; Szymanski, S. K.; Gill, K. S.; Procko, E. Mapping Interaction Sites on Human Chemokine Receptors by Deep Mutational Scanning. J. Immunol. 2018, 200, 3825–3839.

(56) Sultan, M. M.; Wayment-Steele, H. K.; Pande, V. S. Transferable Neural Networks for Enhanced Sampling of Protein Dynamics. J. Chem. Theo. Comput. 2018, 14, 1887–1894.

(57) Moffett, A. S.; Shukla, D. On the transferability of time-lagged independent components between similar molecular dynamics systems. arXiv:1710.00443 2017,

