## Supplementary Methods and Results for "TLmutation: predicting the effects of mutations using transfer learning"

### Supporting information for TLmutation: predicting the effects of mutations using transfer learning.

#### Maximization of the correlation between the predicted values of experiments and the actual experimental values along $w$ and $w'$.

As explained in the main text, We introduced a new Potts model to predict the function with the following modified potentials:

$$J_{ij}^{func.} = w_{ij}.J_{ij} \tag{1}$$

$$h_i^{func.} = w_i'.h_i \tag{2}$$

where $w$ and $w'$ were binary weight matrices ($w_c \in \{0, 1\}$). The values of $w$ and $w'$ are calculated by maximizing the correlation between the predicted values of the experiments and the actual experimental values in the training set ($\hat{Y}^{tr}$ and $Y^{tr}$).

$$\max_{w,w'} \mid \rho(rg_{\hat{Y}^{tr}}, rg_{Y^{tr}}) \mid \tag{3}$$

In another word we have to find subsets of $w$ and $w'$ that maximizes the correlation. For a protein with $n$ amino acids, $w$ will have $\frac{(n-1)(n-2)}{2}$ elements and $w'$ will have $n$. Therefore, the optimization space will have $2^{\frac{(n-1)(n-2)}{2}+n}$ possibilities, which is equal to $6 \times 10^{29}$ for a small protein of size 100. Since the optimization space is so big, we break down the optimization into two steps in series, first optimizing over $w$ and then over $w'$. Each step itself involves three smaller steps. For example for optimizing over $w$, first we test the effect of each element of $w$ ($w_i$) on $\rho(rg_{\hat{Y}^{tr}}, rg_{Y^{tr}})$ by nullifying every element other than $w_i$. Second, we make a list of all $w_i$s and their corresponding values of $\rho(rg_{\hat{Y}^{tr}}, rg_{Y^{tr}})$ sorted based on $\rho(rg_{\hat{Y}^{tr}}, rg_{Y^{tr}})$, which shows how did each $w_i$ affect the correlation in the first step. In the third step, we want to test the accumulated effect of $w_i$s. For all $w_i$s We put $w_i = 0$, then starting from the first $w_i$ in the sorted list, we turn $w_i$ "ON" ($w_i = 1$), calculate $\rho(rg_{\hat{Y}^{tr}}, rg_{Y^{tr}})$, and continue adding $w_i$s to the last element in the list. Keeping tack of the values of $\rho(rg_{\hat{Y}^{tr}}, rg_{Y^{tr}})$ in the third step, we can find the maximum value and the corresponding set of "ON" $w_i$s. We go though the sames steps again for $w'$.

Table S1: **Mutagensis datasets used in case study 1.**

| **Protein** | **UniprotID** | **# Mutations** | **PMID** |
|---|---|---|---|
| HIV capsid protein p24[1] | POL_HV1N5 | 38 | 25102049 |
| DNA Methylase HaeIII[2] | MTH3_HAEAE | 1957 | 26274323 |
| Hepatitis C NS5A[3] | POLG_HCVJF | 1632 | 24722365 |
| Kanamycin kinase[4] | KKA2_KLEPN | 5017 | 24914046 |
| $\beta$-glucosidase[5] | (Sequence from dataset) | 3000 | 26040002 |
| PABP (RRM domain)[6] | PABP_YEAST | 1187 | 24064791 |
| YAP1 (WW domain)[7] | YAP1_HUMAN | 319 | 23035249 |
| Gal4 (DNA-binding domain)[8] | GAL4_YEAST | 1195 | 25559584 |
| PSD95 (PDZ domain)[9] | DLG4_RAT | 1578 | 23041932 |
| Influenza hemaglutinin[10] | (Sequence from paper) | 10721 | 27271655 |
| HSP90 (ATPase domain)[11] | HSP82_YEAST | 4324 | 27068472 |
| Photoactivase yellow protein[12] | PYP_HALHA | 126 | 20889915 |
| UBE4B (U-box domain)[13] | UBE4B_MOUSE | 900 | 23509263 |
| Ubiquitin[14] | RL401_YEAST | 1366 | 24862281 |
| GB1 domain[15] | (Sequence from dataset) | 1045 | 25455030 |
| BRCA1 (Ring domain)[16] | BRCA1_HUMAN | 4873 | 25823446 |
| $\beta$-lactamase[17] | BLAT_ECOLX | 4997 | 25723163 |

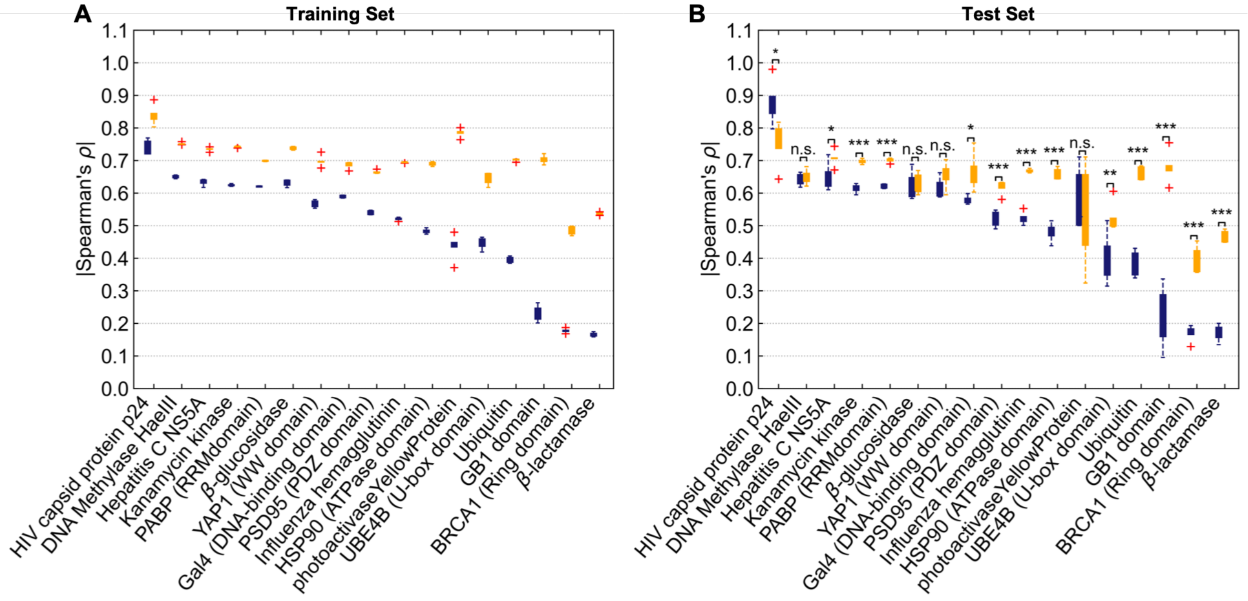


Figure S1: Spearman scores of 17 systems in case study 1 on their (**A**) training and (**B**) test set with random training/test split. Asterisks indicate statistically significant (*, $p < 0.05$; **, $p < 0.01$; ***, $p < 0.001$, *n.s.*, not significant).

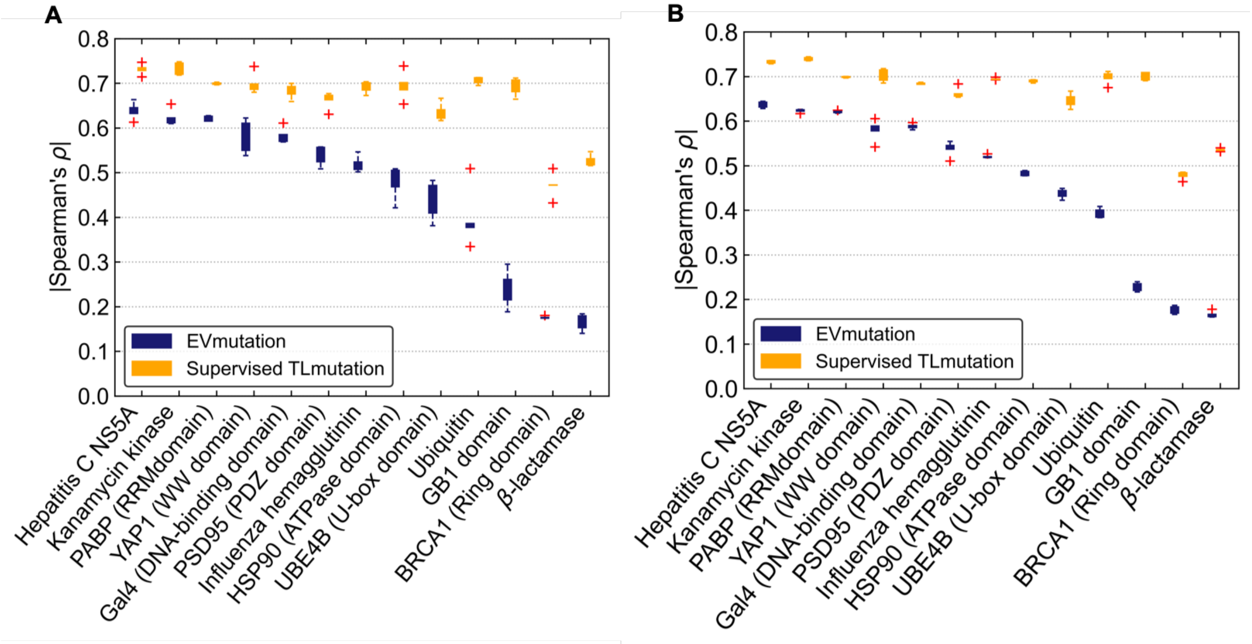


Figure S2: Spearman scores of 13 systems in case study 1 on their training set with training/test split based on (**A**) sequence sites and (**B**) substituted amino acids.

Table S2: **Mutagensis datasets used in case study 2.**

| **Protein** | **UniprotID** | **# Single point mutations** | **# Double point mutations** | **PMID** |
|---|---|---|---|---|
| Trypsin[18] | TRY2_RAT | 14 | 8 | 19703402 |
| FYN[19] | FYN_HUMAN | 41 | 6 | 14556750 |
| PABP (RRM domain)[6] | PABP_YEAST | 1187 | 36522 | 24064791 |
| GB1 domain[15] | (Sequence from dataset) | 1045 | 509693 | 25455030 |

#### Deep mutational scanning dataset of Heredia *et al.*

We will briefly describe the methodology of deep mutational scanning and biochemical experiments used to obtain CXCR4 and CCR5 datasets; however, the original manuscript provides a more comprehensive outline of the procedures.[20]

First, a library of DNA sequences, consisting of the sequences of every possible amino acid substitution, is constructed. CXCR4 and CCR5 single-site saturation mutagenesis (SSM) libraries consisted of >99% of possible amino acid substitutions. Each DNA sequence is transfected into individual cells, in which one cell will express one mutant protein. Cells are stained, typically with antibodies, in order to be detected by fluorescence or microscopy. To detect surface expression, cells are stained with a fluorescent antibody recognizing a small detection tag strategically placed on the protein. BiFC assay detects protein-protein interactions by fluorescence, and can be used to assess how two chemokine receptors bind together to make a functional complex. Fluorescent cells are then sorted by fluorescence-activated cell sorting (FACS), which individually sorts cells based on emitted fluorescence level. Once sorted, the RNA is extracted from cells and sequenced. Mutational datasets for CXCR4 and CCR5 were obtained from the National Center for Biotechnology Information's Gene Expression Omnibus.

### References


(1) Mann, J. K.; Barton, J. P.; Ferguson, A. L.; Omarjee, S.; Walker, B. D.; Chakraborty, A.; Ndung'u, T. The Fitness Landscape of HIV-1 Gag: Advanced Modeling Approaches and Validation of Model Predictions by In Vitro Testing. *PLoS Comput. Biol.* **2014**, *10*, e1003776.

(2) Rockah-Shmuel, L.; Tóth-Petróczy, Á.; Tawfik, D. S. Systematic Mapping of Protein Mutational Space by Prolonged Drift Reveals the Deleterious Effects of Seemingly Neutral Mutations. *PLoS Comput. Biol.* **2015**, *11*, e1004421.

(3) Qi, H. et al. A Quantitative High-Resolution Genetic Profile Rapidly Identifies Sequence Determinants of Hepatitis C Viral Fitness and Drug Sensitivity. *PLOS Pathog.* **2014**, *10*, e1004064.

(4) Melnikov, A.; Rogov, P.; Wang, L.; Gnirke, A.; Mikkelsen, T. S. Comprehensive mutational scanning of a kinasein vivoreveals substrate-dependent fitness landscapes. *Nucleic Acids Res.* **2014**, *42*, e112–e112.

(5) Romero, P. A.; Tran, T. M.; Abate, A. R. Dissecting enzyme function with microfluidic-based deep mutational scanning. *Proc. Natl. Acad. Sci. U.S.A.* **2015**, *112*, 7159–7164.

(6) Melamed, D.; Young, D. L.; Gamble, C. E.; Miller, C. R.; Fields, S. Deep mutational scanning of an RRM domain of the Saccharomyces cerevisiae poly(A)-binding protein. *RNA* **2013**, *19*, 1537–1551.

(7) Araya, C. L.; Fowler, D. M.; Chen, W.; Muniez, I.; Kelly, J. W.; Fields, S. A fundamental protein property, thermodynamic stability, revealed solely from large-scale measurements of protein function. *Proc. Natl. Acad. Sci. U.S.A.* **2012**, *109*, 16858–16863.

(8) Kitzman, J. O.; Starita, L. M.; Lo, R. S.; Fields, S.; Shendure, J. Massively parallel single-amino-acid mutagenesis. *Nat. Methods* **2015**, *12*, 203–206.

(9) Jr, R. N. M.; Poelwijk, F. J.; Raman, A.; Gosal, W. S.; Ranganathan, R. The spatial architecture of protein function and adaptation. *Nature* **2012**, *491*, 138–142.

(10) Doud, M.; Bloom, J. Accurate Measurement of the Effects of All Amino-Acid Mutations on Influenza Hemagglutinin. *Viruses* **2016**, *8*, 155.

(11) Mishra, P.; Flynn, J. M.; Starr, T. N.; Bolon, D. N. Systematic Mutant Analyses Elucidate General and Client-Specific Aspects of Hsp90 Function. *Cell. Rep.* **2016**, *15*, 588–598.

(12) Philip, A. F.; Kumauchi, M.; Hoff, W. D. Robustness and evolvability in the functional anatomy of a PER-ARNT-SIM (PAS) domain. *Proc. Natl. Acad. Sci. U.S.A.* **2010**, *107*, 17986–17991.

(13) Starita, L. M.; Pruneda, J. N.; Lo, R. S.; Fowler, D. M.; Kim, H. J.; Hiatt, J. B.; Shendure, J.; Brzovic, P. S.; Fields, S.; Klevit, R. E. Activity-enhancing mutations in an E3 ubiquitin ligase identified by high-throughput mutagenesis. *Proc. Natl. Acad. Sci. U.S.A.* **2013**, *110*, E1263–E1272.

(14) Roscoe, B. P.; Bolon, D. N. Systematic Exploration of Ubiquitin Sequence, E1 Activation Efficiency, and Experimental Fitness in Yeast. *J. Mol. Biol.* **2014**, *426*, 2854–2870.

(15) Olson, C. A.; Wu, N. C.; Sun, R. A comprehensive biophysical description of pairwise epistasis throughout an entire protein domain. *Curr. Biol.* **2014**, *24*, 2643–2651.

(16) Starita, L. M.; Young, D. L.; Islam, M.; Kitzman, J. O.; Gullingsrud, J.; Hause, R. J.; Fowler, D. M.; Parvin, J. D.; Shendure, J.; Fields, S. Massively Parallel Functional Analysis of BRCA1 RING Domain Variants. *Genetics* **2015**, *200*, 413–422.

(17) Stiffler, M. A.; Hekstra, D. R.; Ranganathan, R. Evolvability as a Function of Purifying Selection in TEM-1 $\beta$-lactamase. *Cell* **2015**, *160*, 882–892.

(18) Halabi, N.; Rivoire, O.; Leibler, S.; Ranganathan, R. Protein Sectors: Evolutionary Units of Three-Dimensional Structure. *Cell* **2009**, *138*, 774–786.

(19) Nardo, A. A. D.; Larson, S. M.; Davidson, A. R. The Relationship Between Conservation, Thermodynamic Stability, and Function in the SH3 Domain Hydrophobic Core. *J. Mol. Biol.* **2003**, *333*, 641–655.

(20) Heredia, J. D.; Park, J.; Brubaker, R. J.; Szymanski, S. K.; Gill, K. S.; Procko, E. Mapping Interaction Sites on Human Chemokine Receptors by Deep Mutational Scanning. *J. Immunol.* **2018**, ji1800343.
